# Simple and cost-effective SNP genotyping method for discriminating subpopulations of the fish pathogen, *Nocardia seriolae*

**DOI:** 10.1101/2021.12.28.474260

**Authors:** Cuong T. Le, Erin P. Price, Derek S. Sarovich, Thu T.A Nguyen, Hung Vu-Khac, Ipek Kurtböke, Wayne Knibb, Shih-Chu Chen, Mohammad Katouli

## Abstract

*Nocardia seriolae* has caused significant fish losses in Asia and the Americas in recent decades, including in Vietnam, which has witnessed devastating economic and social impacts due to this bacterial pathogen. Surveillance strategies are urgently needed to mitigate *N. seriolae* dissemination in Vietnamese aquaculture and mariculture industries. Whole-genome sequencing (WGS) offers the highest level of resolution to discriminate closely related strains and to determine their putative origin and transmission routes. However, WGS is impractical for epidemiological investigations and pathogen surveillance due to its time-consuming and costly nature, putting this technology out-of-reach for many industry end-users. To overcome this issue, we targeted two previously characterised, phylogenetically informative single-nucleotide polymorphisms (SNPs) in *N. seriolae* that accurately distinguish: i) Vietnamese from non-Vietnamese strains, and ii) the two Vietnamese subclades. Using the mismatch amplification mutation assay (MAMA) format, we developed assays that genotype strains based on differences in amplicon melting temperature (melt-MAMA) and size (agarose-MAMA). Our MAMA assays accurately genotyped strains both from culture and fish tissues at low cost, using either real-time (~AUD$1/per sample) or conventional (~AUD$0.50/per sample) PCR instrumentation. Our novel assays provide a rapid, reproducible, and cost-effective tool for routine genotyping of this pathogen, allowing faster identification and treatment of nocardiosis-effected permit fish within Vietnamese aquaculture/mariculture facilities, an essential step in mitigating *N. seriolae*-associated losses.

## Introduction

*Nocardia seriolae*, the aetiological agent of granulomatous disease (nocardiosis) in fresh and marine fish species, has recently become one of the major emerging pathogens impacting global aquaculture (Labrie et al., 2008, Maekawa et al., 2018). *N. seriolae* infection can present as either chronic or acute disease, resulting in up to 70% cumulative mortality. This bacterial pathogen was first reported in farmed yellowtail in Japan in 1967 (Kariya et al., 1968), and has subsequently been documented in more than 50 fish species in Taiwan, China, Vietnam, Korea, USA, and Mexico, where it has been associated with considerable economic losses (Chen et al., 1989, Chen and Tung, 1991, Kudo et al., 1988, Chen et al., 2000, Huang, 2004, Park et al., 2005, Shimahara et al., 2008, Shimahara et al., 2009, Cornwell et al., 2011, Vu-Khac et al., 2016, Del Rio-Rodriguez RE, 2021).

Existing methods for characterising *N. seriolae* strains have demonstrated weaknesses that have impeded effective infection prevention and control of nocardiosis in fish. Since *N. seriolae* is highly clonal (Han et al., 2018, Le et al., 2021), differences in α-glucosidase activity, drug susceptibility, and biased sinusoidal field gel electrophoresis and pulsed-field gel electrophoresis (PFGE) profiles are often insufficient resolution to accurately determine relationships among isolates (Shimahara, 2006, Shimahara et al., 2008, Shimahara et al., 2009). In addition, it is challenging to accurately compare electrophoresis profiles between laboratories (Abdelbary et al., 2017, Krawczyk and Kur, 2018). These limitations have made it difficult to understand the epidemiological, temporal and geographic variation of *N. seriolae* collected from different countries (Le et al., 2021).

Whole-genome sequencing (WGS) is becoming a popular tool for epidemiologic analysis of bacterial pathogens (Uelze et al., 2020). By identifying single-nucleotide polymorphism (SNP) variation across the genome and phylogenetically reconstructing population diversity using these genome-wide SNPs, WGS yields very high resolution that enables the discrimination of closely related strains, an important facet in accurate source tracing efforts (Schürch et al., 2018). Although WGS is considered the ‘gold standard’ for genotyping bacterial pathogens, this method is currently out-of-reach for lower resourced regions or for industry end-users that lack access to specialised molecular equipment and computational capacity (Mitchell et al., 2017).

Cheap, rapid, simple, and accessible SNP genotyping methods offer an attractive alternative to WGS for characterising bacterial pathogens. By targeting phylogenetically informative SNPs, these methods can offer similar levels of epidemiologically useful information, but at much lower cost and labour-intensity than WGS. Several high-throughput SNP genotyping techniques have been developed using fluorogenic probes e.g. TaqMan, Amplifluor, and SNP arrays (Raviv et al., 2008, Birdsell et al., 2012, Zhang et al., 2015). However, these techniques are expensive and require specialised equipment, making them impractical for routine isolate genotyping, particularly in lower-resourced settings (Mitchell et al., 2017). There is a need for simple, accurate, and cost-effective genotyping assays for characterising *N. seriolae* isolates that can be implemented in laboratories lacking access to sophisticated equipment.

SNP interrogation using the mismatch amplification mutation assay (MAMA) format has been used successfully to genotype several pathogens, including *Francisella tularensis, Burkholderia pseudomallei, B. mallei* (Birdsell et al., 2014), *Yersinia pestis* (Mitchell et al., 2017), *Neisseria gonorrhoeae* (Donà et al., 2018), *Bacillus anthracis* (Price et al., 2010, Lekota et al., 2020), *Mycoplasma synoviae* (Kreizinger et al., 2015), *Mycobacterium avium* (Leão et al., 2016), *Mycoplasma gallisepticum* (Sulyok et al., 2019), and *Mycoplasma hyopneumoniae* (Felde et al., 2020). For *N. seriolae*, we have previously developed SYBR green-based assays (SYBR-MAMA) to differentiate strains from infected fish propagated in Vietnamese mariculture facilities (Le et al., 2021). Shortcomings of the SYBR-MAMA approach are that two mastermixes are required for each assay and for each strain, and only the real-time PCR platform can be used, which reduces its cost-effectiveness and accessibility. Here, we examined single-tube MAMA formats (melt-MAMA and agarose-MAMA) to determine whether these methods could perform cost-effective *N. seriolae* SNP genotyping using both real-time and conventional PCR equipment. We demonstrate the feasibility of *N. seriolae* genotyping using these single-tube MAMA formats by testing them on DNA obtained directly from cultures and from infected fish tissues.

## Methods

### Ethics statement

No ethics approval was required for harvesting kidneys from dead farmed fish specimens, which were obtained directly from Vietnamese fish farmers using strict hygiene measures. Harvesting of kidneys from *N. seriolae*-challenged permit fish (*Trachinotus falcatus*) with VT_45 was approved by the University of the Sunshine Coast (USC) Animal Ethics Committee (approval no. ANS1861). The study did not involve endangered or protected animals.

### DNA extraction from bacterial cultures and fish tissues

Total genomic DNA (gDNA) was obtained from 60 *N. seriolae* strains and 15 kidney samples of permit fish collected from a challenge experiment (*n*=5) and dead permit fish (*Trachinotus falcatus)* from a farm (*n*=10). Due to a ban on *N. seriolae* importation into Australia, all live culture and infected fish tissue work were carried out in Vietnam and Taiwan, with DNA confirmed to be sterile prior to importation. RNAlater or 90% ethanol-fixed fish kidney samples were imported to USC for further testing.

Of the 60 strains, nine were isolated between 2003 and 2007 from six different fish in Taiwan and were included as “outgroup” (i.e. non-Vietnamese) strains. The remaining 51 strains were isolated from farmed *T. falcatus* in four Vietnamese provinces (Phú Yên, Khánh Hòa, Ninh Thuận, and Vũng Tàu) in the South Central Coast region in 2014 and 2015 (Table 1). Seven of these strains have previously been genome-sequenced, with strains KH_11, PY_31, VT_45 belonging to Vietnam Clade 1 while KH_21, PY_37, NT_50, VT_62 belong to Vietnam Clade 2 according to phylogenomic analysis (Le et al., 2021). DNA from these two clades were used as positive controls for the melt-MAMA and agarose-MAMA assays developed in this study.

**Table 1.**
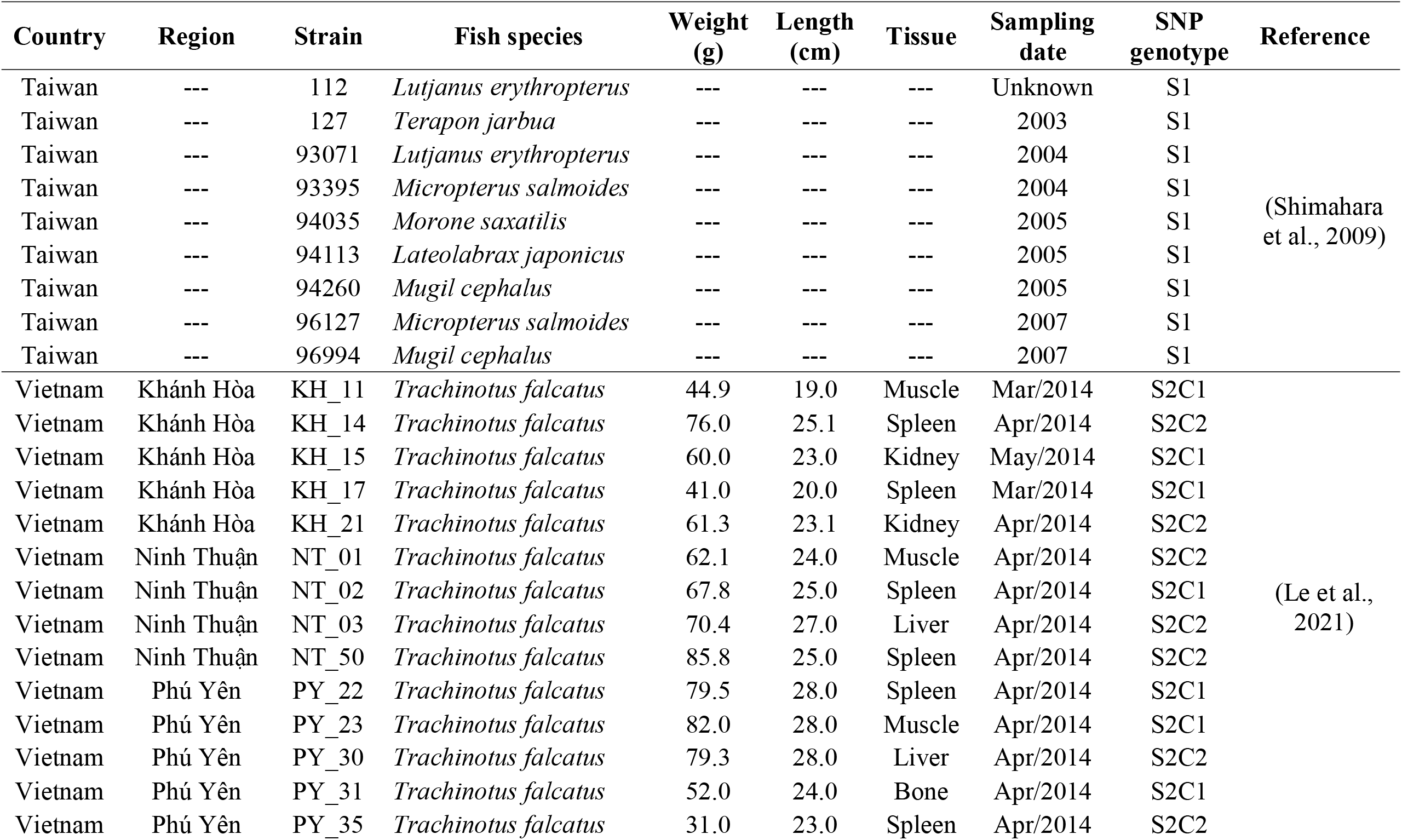
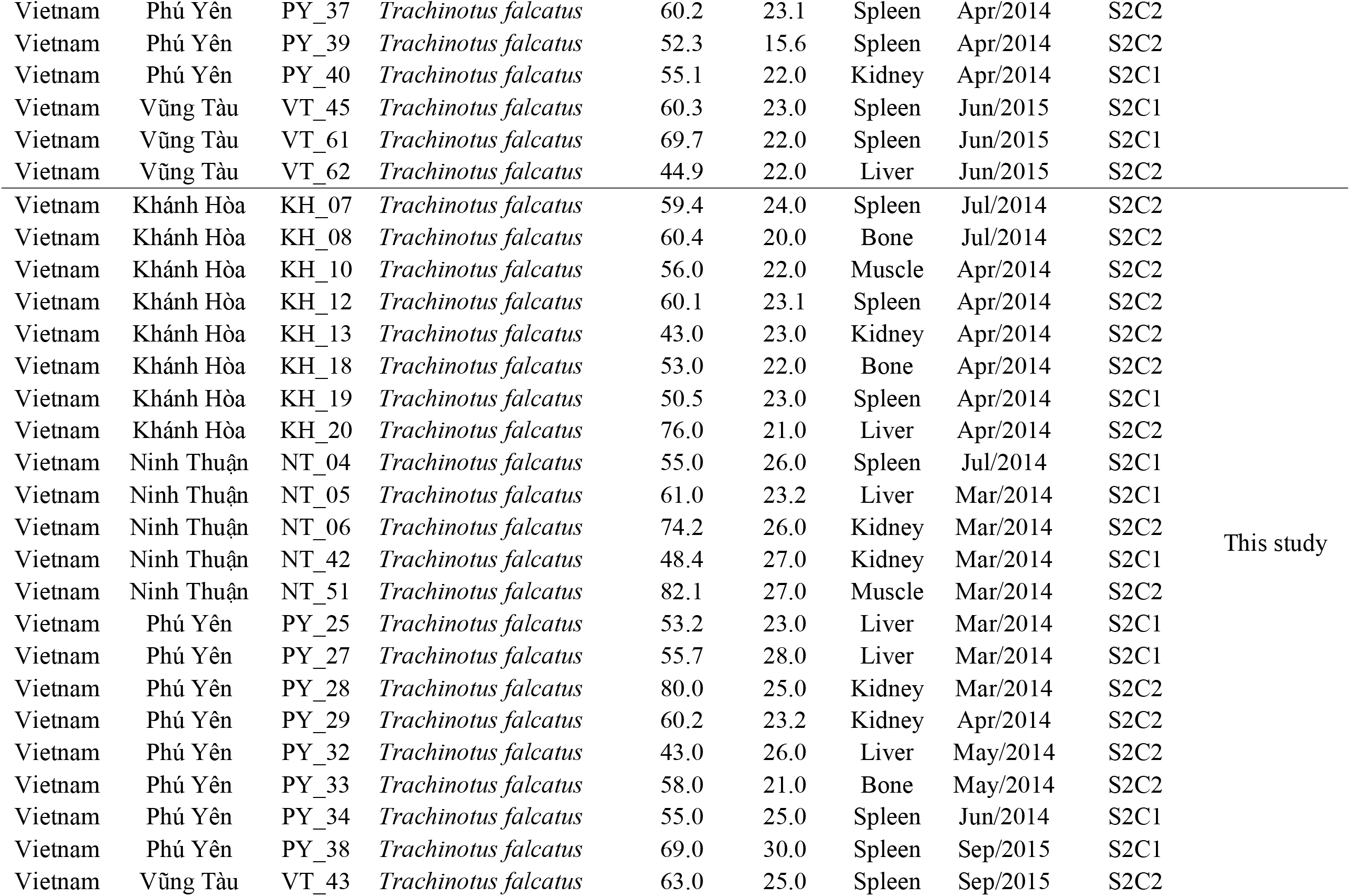
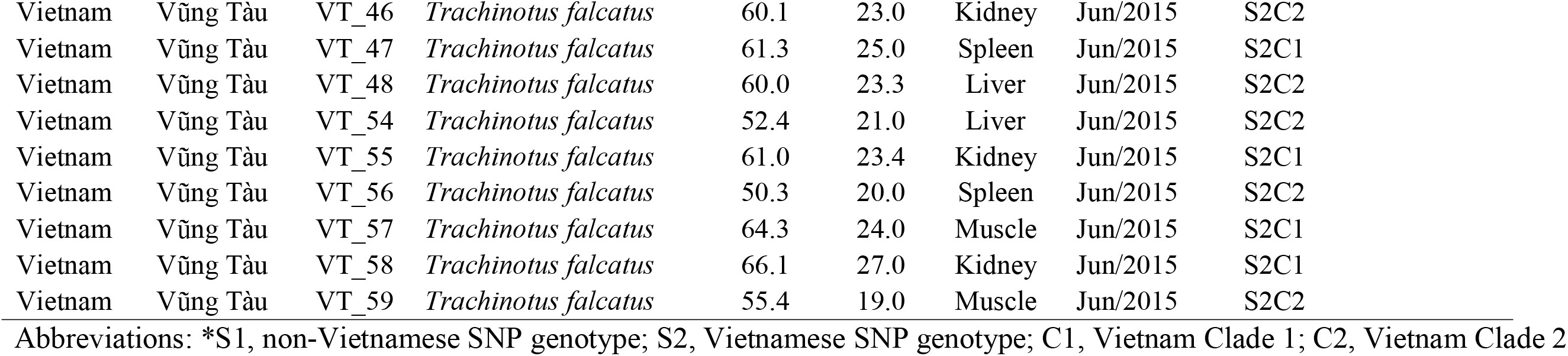
*Nocardia seriolae* strains collected in this study, and their single-nucleotide polymorphism (SNP) genotype profiles

*N. seriolae* gDNA was extracted from 5-day old cultures using the Wizard® Genomic DNA Purification Kit (Promega, USA) as per the manufacturer’s instructions. For gDNA of kidney samples, tissues were incubated until dissolved at 37°C in 495 μL extraction buffer [(4 M urea, 0.2 M NaCl, 1 mM trisodium citrate, 1% SDS (Sigma-Aldrich)] supplemented with 5 μL Proteinase K (20 mg mL^−1^; Bioline Australia) to lyse cells. Protein, cellular debris, and detergent were removed by centrifugation in 7.5 M ammonium acetate at 14000×*g* for 3 min at 4°C. Nucleic acids were recovered by isopropanol precipitation at 14000×*g* for 10 min. The nucleic acid pellet was washed twice with ethanol and eluted in 100 μL water containing 10 μM Tris-HCl and 0.05% Triton X-100 (v/v; Sigma-Aldrich). DNA quantity and purity were assessed using a NanoDrop 2000 (Thermo Scientific, Scoresby, VIC, Australia).

### SNP selection

We have previously shown that biallelic SNPs at positions 60409 and 587171 in the EM150506 reference genome (Han et al., 2018) are phylogenetically informative for differentiating: i) Vietnamese strains from non-Vietnamese strains, and: ii) the two Vietnamese clades, respectively (Le et al., 2021). We targeted these same SNPs for melt- and agarose-MAMA development.

### Assay design

MAMA design typically involves using two allele-specific (AS) primers and a single common primer to differentiate a biallelic SNP of interest. Melt-MAMA and agarose-MAMA differ from SYBR-MAMA in that all primers are included together in one tube, rather than across two separate AS primer reactions (Birdsell et al., 2012). The AS primers in melt-MAMA and agarose-MAMA compete for the same SNP locus on the DNA template, with the 3’ matched template being much more robust at binding and extending its product, essentially drowning out any signal that might be generated from the mismatched AS primer. In melt-MAMA, the difference in melting temperature (*T*_m_) is detected by using SYBR Green and post-PCR melt curve analysis on the real-time PCR platform, whereas in agarose-MAMA, the difference in amplicon size is detected by 3% agarose gel electrophoresis (Birdsell et al., 2012, Mitchell et al., 2017).

Most primers used for our melt- and agarose-MAMAs have been previously published in SYBR-MAMA format (Le et al., 2021). The exceptions were CtS1_nonViet_For2 and CtS2_Clade2_Rev2, which included a modified 5’ 20bp-long GC-rich clamp (ggggcggggcggggcggggc) compared with CtS1_nonViet_For and CtS2_Clade2_Rev (Le et al., 2021) to increase the *T*_m_ and size of the corresponding amplicon (Birdsell et al., 2012). These new primers also included different penultimate mismatches at the 3’-end to the previously published primers. This difference creates two mismatched nucleotides in the 3’ region of the primer for the non-allelic template, but only one difference in the correct allele template. Allele-specific primers carrying a single mismatch has no effect on the overall PCR yield, whereas primers carrying 2 consecutive mismatches fail to generate any detectable amplification products (Fig. 1). Genome location, primer sequences, and amplicon sizes for the MAMAs developed in this study are listed in Table 2.

**Fig. 1.**
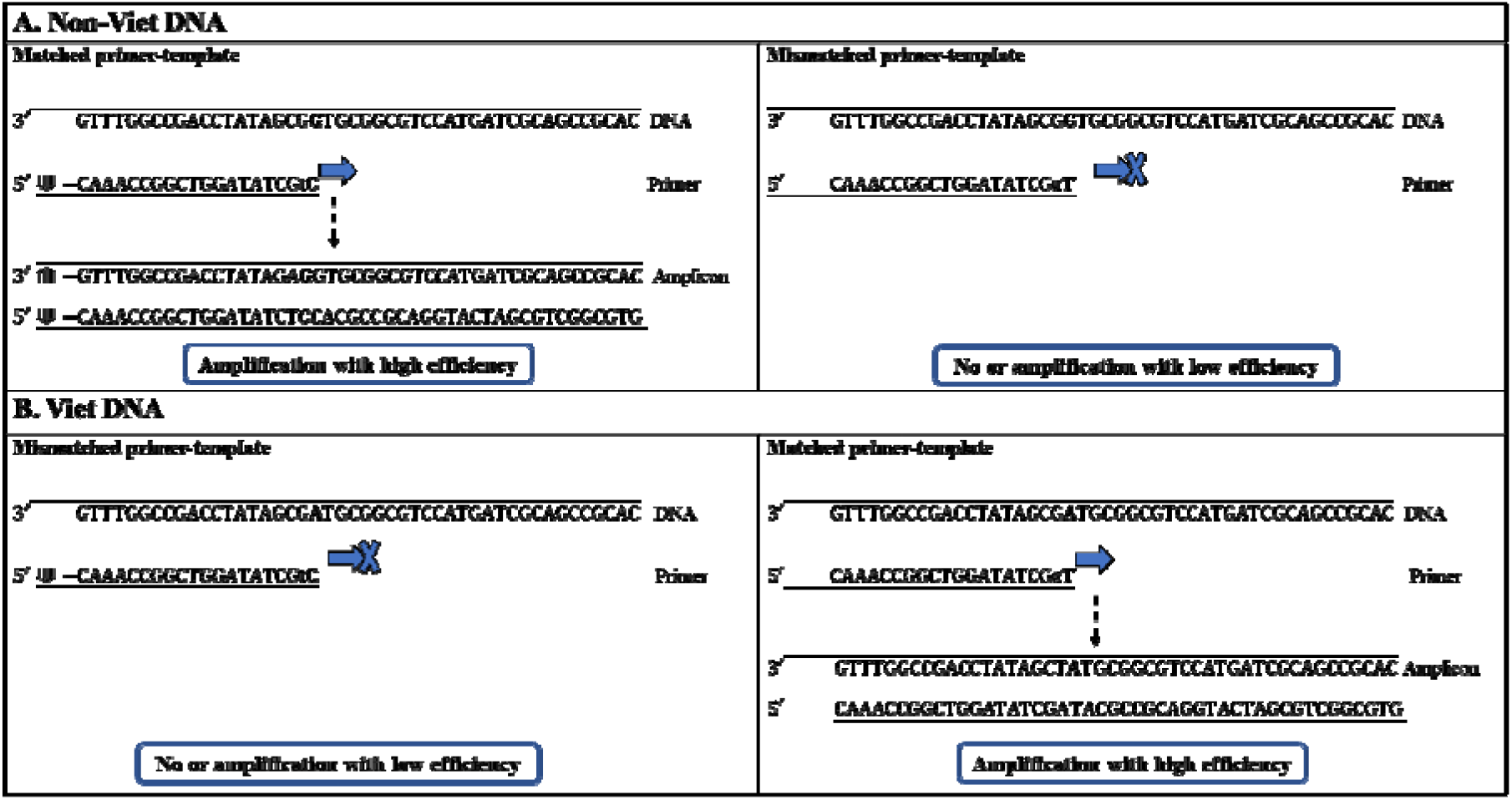
Schematic representation of mismatch amplification mutation assay (MAMA) PCR reaction for discriminating Viet and Non-Viet *Nocardia seriolae* strains. Two allele-specific (AS) primers compete for the same SNP locus on either non-Viet (A) or Viet (B) DNA template with non-Viet primer labelled with GC clamp (⟱). Taq Polymerase (blue arrow) extends from the 3’ matched AS primer that has single mismatch to the template but fail to do that (blue arrow with X) for the primer with two mismatches. This indicates the amplification of the perfect-matched amplicon and little to no amplification of the mismatched amplicon. The GC –clamp ‘‘labeled’’ amplicons are larger than non-GC amplicons in size.

**Table 2.**
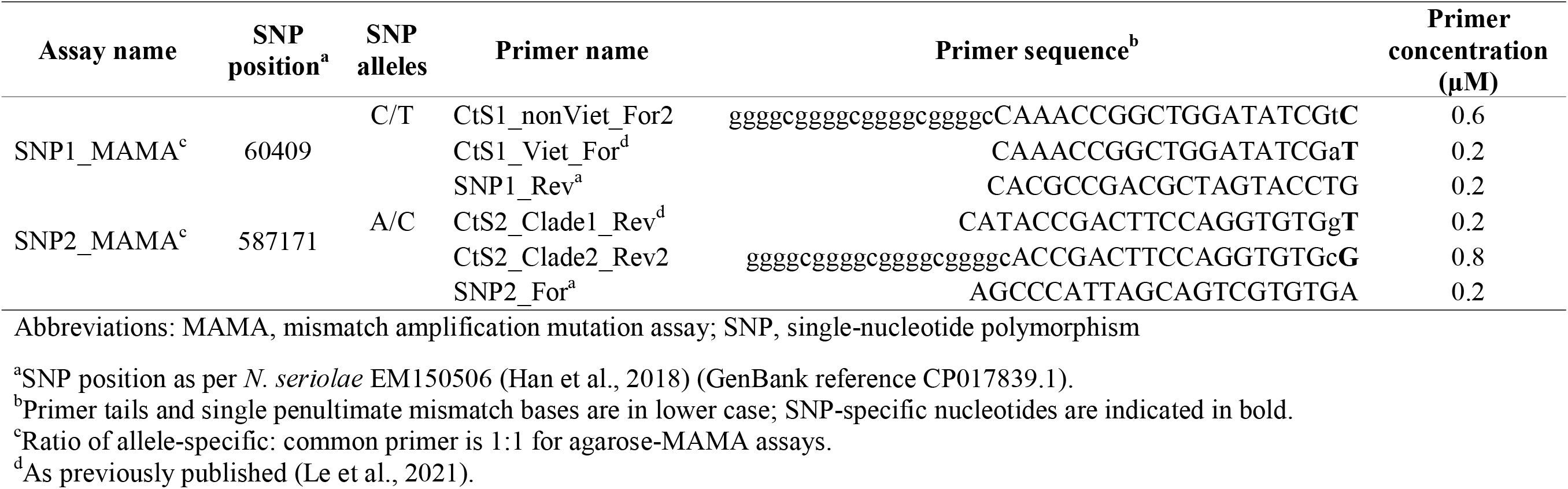
Primer sequences of melt-MAMA and agarose-MAMA assays for differentiating Vietnamese *Nocardia seriolae* strains.

**Table 3.** Reference values for mismatch amplification mutation assays (MAMAs) based on *Nocardia seriolae* control isolates

| Strain/s | Genotype | Melt-MAMA |  |  |  | Agarose-MAMA |  |
| --- | --- | --- | --- | --- | --- | --- | --- |
|  |  | SNP1 C<sub>T</sub> | SNP1 T<sub>m</sub> | SNP2 C<sub>T</sub> | SNP2 T<sub>m</sub> | SNP1 product size (bp) | SNP2 product size (bp) |
|  | Non_Viet | 32.2 ± 0.5 | 86.7 ± 0.0 | --- | --- | 66 | --- |
|  | Viet | 34.1 ± 0.4 | 79.4 ± 0.4 | --- | --- | 46 | --- |
|  | Viet_Clade1 | --- | --- | 28.9 ± 0.7 | 80.2 ± 0.2 | --- | 48 |
|  | Viet_Clade2 | --- | --- | 29.1 ± 0.4 | 88.8 ± 0.0 | --- | 65 |

Abbreviations: bp, base pairs; C<sub>T</sub>, cycles-to-threshold (value); MAMA, mismatch amplification mutation assay; T<sub>m</sub>, melting temperature; SNP, single- nucleotide polymorphism

### Melt-MAMA and agarose-MAMA PCRs

To assess SNP genotyping accuracy with the newly designed melt-MAMA and agarose-MAMA assays, PCRs were performed using the Taiwanese and Vietnamese control DNA samples. Once each assay was optimised, the remaining 51 Vietnamese isolates were screened using both formats. Nuclease-free water was used as a negative control in all PCR assays. All samples were run in duplicate or at higher replication when deemed necessary.

All melt-MAMA PCRs were performed on a CFX96 Real-Time PCR Detection System using white hard shell 96-well PCR plates or 0.2 mL 8-tube PCR strips (Bio-Rad, Gladesville, NSW, Australia), and PCR results examined using the CFX Maestro v4.1.2433.1219 software (Bio-Rad). Melt-MAMAs were performed in 10 μL total reaction volume, containing 1 ng of target DNA template (isolate or fish tissue DNA), 1X SsoAdvanced™ Universal SYBR® Green Supermix (Bio-Rad), primers (Macrogen, Inc., Geumcheon-gu, Seoul, Republic of Korea) at concentrations shown in Table 2, and RNase/DNase-free PCR-grade water (Thermo Fisher Scientific). Thermocycling conditions comprised an initial 2 min denaturation at 95°C, followed by 40 cycles of 95°C for 15 sec and 60°C for 15 sec. After amplification, a melting curve dissociation analysis was performed comprising 95°C for 15 sec, followed by incremental temperature ramping (0.3°C) from 60°C to 95°C. *T*_m_ values were determined by visual inspection of the melting curves. The assays were optimised by altering AS primer ratios, as previously described (Birdsell et al., 2012).

### Agarose-MAMA assays

Agarose-MAMAs were carried out in a 25 μL total volume, containing 1 μL of target DNA template at ~10ng/μL, primers (Table 2), and 1X AmpliTaq Gold 360 master mix (Applied Biosystems, Foster City, CA, USA). Cycle conditions were: 94°C for 5 min, followed by 40 cycles of 95°C for 30 sec, 60°C for 30 sec, and 72°C for 30 sec, followed by a final extension at 72°C for 5 min. All amplified PCR products were electrophoretically analyzed in 3% agarose gel (MetaPhor Agarose, Lonza Group) stained with 0.5 μg/mL ethidium bromide. The 100-bp DNA ladder (Axygen) was used for size referencing.

### Assay validation

To assess assay capability to genotype *N. seriolae* directly from DNA samples of fish kidneys, we tested the performance of each assay on positive control samples obtained from the challenge experiment with our positive control *N. seriolae* DNA extract (strain VT_45). To further assess this capability, we tested the assays on 10 kidney DNA samples confirmed to be *N. seriolae* positive from a permit fish farms in Nha Trang city by our newly designed *N. seriolae*-specific-TaqMan-assay.

## Results

Melt-MAMA and agarose-MAMA testing of our control samples, and subsequently our test samples, demonstrated 100% sensitivity, with successful differentiation of target and non-target strains for all assays (Fig.1 and Fig.2) at the same accuracy as WGS, as previously reported (Le et al., 2021). Negative controls did not amplify, nor generated nonspecific products with melt profiles differing from the profiles of the expected two melting temperatures and sizes of target amplicons.

**Fig. 2.**
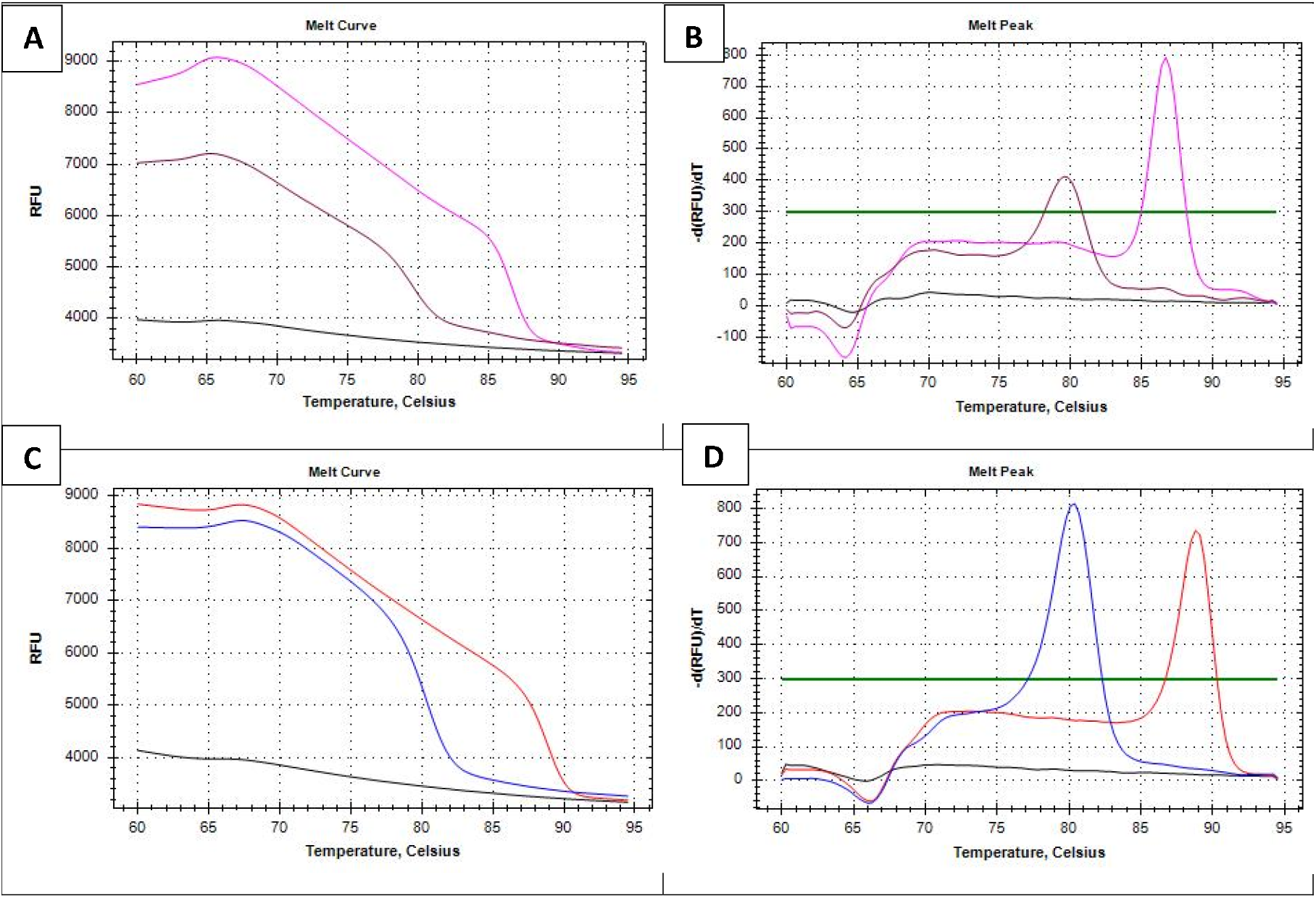
Raw (left) and derivative (right) melt-MAMA melt curves for single-nucleotide polymorphism (SNP) discrimination of *Nocardia seriolae* strains. **Panels A and B**. SNP1 assay results for Viet (maroon line, melting temperature [*T*_m_]: 80.3°C) and non-Viet (pink line, *T*_m_: 86.8°C) strains. **Panels C and D**. SNP2 assay results for Viet_Clade1 (blue line, *T*_m_: 80.3°C) and Viet_Clade2 (pink line, *T*_m_: 88.5°C) strains. Black lines represent negative controls.

Using either MAMA formats, the C→T polymorphism at locus 60409 (SNP1) was clearly identified, with the 51 Vietnamese strains readily distinguished from the nine Taiwanese *N. seriolae* strains. Due to the 5’GC-tail on the non_Viet AS primer, the melt-MAMA *T*_m_ for this genotype (86.7°C ± 0.0) is ~6.5°C higher than for the Viet genotype (79.4°C ± 0.4) (Fig.1 and Fig.2). Similarly, the agarose-MAMA amplicon size for the non_Viet *N. seriolae* genotype (66bp) is 20bp longer than the Viet genotype (46bp) and can be readily distinguished on a 3% agarose gel (Fig.3). As with SNP1, the A→C polymorphism at locus 587171 (SNP2) was clearly identified among the 51 Vietnamese isolates, with an ~8.6°C *T*_m_ difference between Viet_Clade1 (80.2 °C ± 0.2) and Viet_Clade2 (88.8°C ± 0.0) strains (Fig.2) using melt-MAMA. Likewise, the agarose-MAMA differentiated the two Viet-clade strain genotypes based on amplicon size, with Viet_Clade1 strains (65bp) yielding a product 17bp longer than Viet_Clade2 strains (48bp) (Fig.3). The melt- and agarose-MAMA profiles for SNP2 were identical, with 22/51 strains identified as Viet_Clade1, and 30/51 strains identified as Viet_Clade2 (Table 1).

**Fig. 3.**
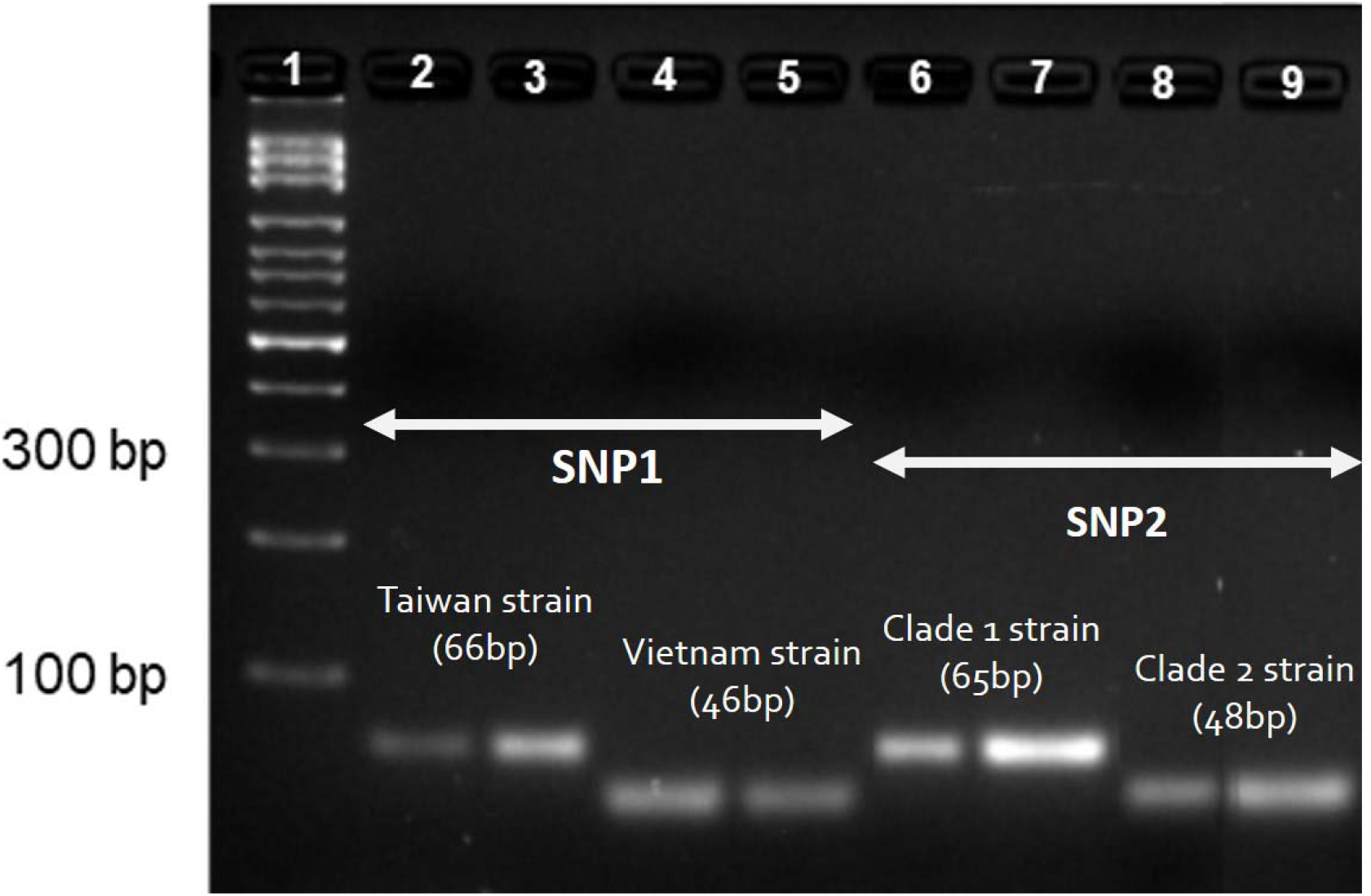
Amplicon size differences in agarose-MAMA single-nucleotide polymorphism (SNP) assays. Lane 1, 100-bp DNA ladder. In the SNP1 assay (Lanes 2-5), non_Viet (Taiwan strains) yield 66bp fragments, whereas Viet strains produce 46bp fragments. In the SNP2 assay (Lanes 6-9), Viet_Clade1 strains yield 65bp fragments, whereas Viet_Clade2 strains produce 48bp fragments.

Following assay validation on cultured isolates, we tested the performance of the melt and agarose-MAMA assays on DNA prepared from fish tissue samples. Melt and agarose-MAMA genotyping of five permit fish kidney samples from an *N. seriolae* challenge experiment confirmed the presence of Viet Clade 1 *N. seriolae* (VT_45) in all five samples, consistent with the strain genotype used for the infection challenge. Next, 8/10 kidney samples from farmed permit fish were melt- and agarose-MAMA-positive; the remaining two samples were below the assay limit of detection. Of these eight positive tissues, all contained the Viet genotype (*n*=8); among these, three were Viet_Clade1, and the remaining five were Viet_Clade2.

Similar to the previous work, we observed non-specific PCR products in a small number of instances. A small number of positive samples generated nonspecific products; however, their profiles were distinct from expected *T*_m_ and amplicon sizes for melt-MAMA and agarose-MAMA, respectively, enabling them to be readily differentiated from ‘real’ genotypes. Previous MAMA studies (Birdsell et al., 2012, Mitchell et al., 2017) have also reported occasional faint, non-specific PCR products; a possible cause of this phenomenon has been suggested to be differences in *Taq* polymerase proofreading activity. Replacement of the MyTaq HS Red mix (Bioline, Eveleigh, NSW, Australia) with AmpliTaq Gold 360 master mix solved the spurious amplification problem in our study. We did not explore different mastermix options for our melt-MAMA assays due to very low fluorescence, and clearly distinct *T*_m_ values for the occasional spurious amplicons.

## Discussion

*N. seriolae* infections have a significant impact on fish farming and food security in Asia, with nocardiosis outbreaks continuing to result in large economic losses due to mass fish mortalities (Labrie et al., 2008, Maekawa et al., 2018, Lei et al., 2020, Liao et al., 2021). Conventional *N. seriolae* genotyping methods e.g. biased sinusoidal field gel electrophoresis, restriction fragment length polymorphism analysis, PFGE) are laborious, time consuming, expensive, low-throughput, and have poor capacity for interlaboratory comparisons (Shimahara et al., 2008, Shimahara et al., 2009, Le et al., 2021). Amplicon sequencing of target genes overcomes these interlaboratory issues but suffers from slow turnaround time and labour intensity. Although considered the ‘gold standard’ genotyping method for bacterial pathogens (Salipante et al., 2015, Uelze et al., 2020), WGS is expensive, slow, complex, labour-intensive, and computationally demanding, and as such remains prohibitive for lower-resourced or commercial settings such as mariculture facilities (Köser et al., 2012, Mitchell et al., 2017, Bayliss et al., 2017). To overcome these shortcomings, we developed novel MAMA genotyping assays targeting two phylogenetically informative SNPs to permit the rapid, simple, and cost-effective identification of Vietnamese strains (SNP1) and their two clades (SNP2). These SNPs were derived from comparative genomic analysis of 19 *N. seriolae* strains, which represent the known genomic diversity of this species (Le et al., 2021). Our MAMAs provide a valuable genotyping tool for epidemiological studies of this bacterium, a matter of importance given the increasing distribution of *N. seriolae* in Vietnamese mariculture areas (Giang et al., 2012, Vu-Khac et al., 2016) and the paucity of inexpensive standardised genotyping schemes for its detection and characterisation (Chen et al., 2000, Han et al., 2018). Our MAMAs yielded identical genotypes to those obtained from WGS (Le et al., 2021), confirming the high accuracy and interlaboratory capacity of this method. Testing of the SNP1 assay across 60 strains – 51 from Vietnam and 9 from Taiwan – yielded 100% concordance with expected genotypes based on geographic origin. Subsequent testing of the SNP2 MAMAs against the 51 Vietnamese strains found approximately equal distribution of Viet_Clade1 and Viet_Clade2 genotypes. This finding confirms our previous work suggesting widespread, unmitigated distribution of these clades throughout Vietnamese mariculture facilities in four South-Central Coast provinces (Khánh Hòa, Ninh Thuận, Phú Yên, and Vũng Tàu) (Le et al., 2021). Most importantly, MAMA removes the need for gene sequencing or WGS, enabling the rapid and inexpensive characterisation of isolates obtained from emerging outbreaks. Our MAMA assays can be performed either on the real-time PCR (SYBR-MAMA or melt-MAMA) or conventional PCR (agarose-MAMA) platform. The melt-MAMA format provides a gel-free, probe-less method that avoids potential contamination issues associated with amplicon handling and costly probe synthesis. For laboratories without real-time instruments or those in a resource poor setting, agarose-MAMA is a viable alternative as its employs conventional PCR coupled with standard agarose gel electrophoresis (Birdsell et al., 2012, Mitchell et al., 2017, Felde et al., 2020).

We have previously developed SYBR-MAMAs to genotype our two *N. seriolae* SNPs (Le et al., 2021); however, this method requires two separate PCRs for each sample to interrogate the alternate allele states, which reduces throughput, increases consumables costs, and doubles the amount of DNA required to interrogate each genotype. In this current study, we made minor changes to our SYBR-MAMA primers to enable single-tube genotyping via melt-MAMA and agarose-MAMA, thereby reducing assay complexity and halving reagent expenditure. We estimate a cost of ~AUD$1/per sample using our melt-MAMAs, ~AUD$0.50/per sample using our agarose-MAMAs. Given their high-throughput capacity, low cost, and simplicity, our MAMAs are practical for large-scale epidemiological investigations. Due to its slow growth, routine isolation and genotyping of *N. seriolae* from infected fish is a fastidious and time-consuming process that has a low success rate (Austin and Austin, 2007, Lewis and Chinabut, 2011, Vu-Khac et al., 2016, Del Rio-Rodriguez RE, 2021). We demonstrate that our melt-MAMA and agarose-MAMA method is capable of genotyping directly from infected fish tissues, with 13/15 tested tissues (including five challenge experiment tissues) able to be genotyped using this method. Agarose-MAMA is therefore a useful surveillance tool for determining the genotype/s of *N. seriolae*-infected fish. Despite showing applicability on tissue samples, the lower limits of our melt- and agarose-MAMAs were not determined in this study. Therefore, further studies are required to assess their detection and quantitation limits, particularly from difficult specimens such as those containing PCR inhibitors (e.g. spleen, blood tissues). In conclusion, our study is the first to apply MAMA to genotype SNPs in a pathogen of aquatic animals. These MAMAs provide a simple, cost-effective, and rapid method for discriminating Vietnamese *N. seriolae* strains from those found in other Asian countries, including directly from diseased fish. Routine implementation of our assays in mariculture surveillance will assist with targeted biocontrol measures (e.g. antibiotic treatment, quarantine), and would mitigate the inadvertent spread of this dangerous infectious disease between areas, particularly in Vietnam, where an estimated 10-20% fish are now chronically infected with this pathogen (Mr. Ut Van Phan, Institute of Aquaculture, Nha Trang university). Our MAMAs will also permit the early detection of *N. seriolae* strains into new geographic regions.

## Author contributions

CL: Project design, sample collection, sample and data analysis, results interpretation, drafting of manuscript.

EPP: Assistance with project design, supervision, data analyses and interpretation, drafting and revising paper.

DSS: Assistance with project design, revising paper.

TTAN: Assistance in the sample collection, preparation and drafting paper.

HV-K: Sample collection guidance, drafting and revising paper.

IK: Assistance with the project design, revising paper.

WK: Supervision, assistance with project design, drafting paper.

S-CC: Providing isolates, drafting paper.

MK: Supervision, project design, funding acquisition, revising paper.

All authors read and approved the final manuscript.

## Conflict of Interest Statement

The authors have no competing interests to declare.

## Acknowledgments

We gratefully acknowledge the financial support and laboratory facilities provided by the Genecology Research Centre, the University of the Sunshine Coast, and Nha Trang University. This research was supported by an Australia Awards PhD scholarship to CL, which is funded by the Australian Department of Foreign Affairs and Trade. DSS and EPP were supported by Advance Queensland fellowships (AQRF13016-17RD2 and AQIRF0362018, respectively).

## Notes

### Competing Interest Statement

The authors have declared no competing interest.

